# Whole Genome Sequencing and Assembly of the House Sparrow, *Passer domesticus*

**DOI:** 10.1101/2023.11.04.565608

**Authors:** Vikas Kumar, Gopesh Sharma, Sankalp Sharma, Samvrutha Prasad, Toral Vaishnoi, Dalia Vishnudasan, Gopinathan Maheswaran, Kaomud Tyagi, Inderjeet Tyagi, Shailesh Desai, PB Kavi Kishor, Gyaneshwer Chaubey, Prashanth Suravajhala

**Affiliations:** Zoological Survey of India, Kolkata, 700053; Bioclues.org, India; NIMS University, Jaipur; Amrita School of Biotechnology, Amrita Vishwa Vidyapeetham, Clappana PO 690525, Kerala, India; Unipath speciality Labs, Ahmedabad, Gujarat; Department of Genetics, Osmania University, Hyderabad 50007, India; Cytogenetics Lab, Department of Zoology, Banaras Hindu University, Varanasi, UP, India; Department of Biosciences, Manipal University Jaipur, Jaipur

**Keywords:** Passerines, house sparrow, next generation sequencing, evolution, adaptation

## Abstract

The common house sparrow, *Passer domesticus* is a small bird belonging to the family Passeridae. Here, we provide high-quality whole genome sequence data along with assembly for the house sparrow. The final genome assembly was assembled using a Shovill/SPAdes/MASURCA/BUSCO workflow, consisting of contigs spanning 268193 bases and coalescing around a 922 MB sized reference genome. We employed rigorous statistical thresholds to check the coverage, as the Passer genome showed considerable similarity to *Gallus gallus* (chicken) and *Taeniopygia guttata* (Zebra finch) genomes, also providing a functional annotation. This new annotated genome assembly will be a valuable resource as a reference for comparative and population genomic analyses of passerine, avian, and vertebrate evolution.

**Significance:** Avian evolution has been of great interest in the context of extinction. Annotating the genomes such as passerines would be of significant interest as we could understand the behavior/foraging traits and further explore their evolutionary landscape. In this work, we provide a full genome sequence of Indian house sparrow, *viz. Passer domesticus* which will serve as a useful resource in understanding the adaptability, evolution, geography, allee effects and circadian rhythms.

## Introduction

Over the last 12 years, a growing number of bird reference genomes have been studied, providing insight into the phylogenetic relationships among the avians [1-9]. The bird genome project [B10K] has been of great significance in providing major scientific breakthroughs in phylogenetics [10,11]. With more than 1200 species, comprising 13% of all known world’s avian, India is blessed with considerable avian diversity. Regrettably, India is ranked 3rd in having rare and threatened avian species in the world [12]. The house Sparrow was introduced to India via Europe from North Africa and Eurasia by the ancient Romans. Sparrows are found in a variety of habitats, including grasslands, forests, deserts, and agricultural areas, as well as in urban areas, such as parks and gardens. They are omnivorous, typically feeding on insects, spiders, worms, seeds, fruits, and grains, and are mainly seed eaters but feed their young ones with insects and other invertebrates; during the breeding periods, they prefer areas with a high abundance of invertebrates[13]. Over the years, there has been a tremendous decline in their population worldwide, and therefore understanding causal mechanisms through which urbanization affects their population remains limited because of numerous perplexing factors, what is believed to be due to rapid urbanization along with deforestation, lack of cavity nesting, along with absence of hedges in modern landscaping. Many hypotheses have been proposed explaining the house sparrow population decline, for example increased predation by domestic cats or sparrow hawks (*Accipiter nisus)*, cleaner streets providing reduced foraging opportunities, competition for food from other urban species, loss of nesting sites, particularly under the eaves and in the roofs of houses, pollution/air quality, both in terms of immediate toxicity and indirect toxicity through the food supply, increased use of pesticides in parks and gardens, disease transmission[14] and finally, the Allee effect [48]. *Passer domesticus*, between 1970-1990 had an extremely large breeding population with over 63 million pairs, and has been declining over the years in Turkey and some parts of the European Union. Analyzing house sparrow habitat characteristics on a fine scale in almost 200 census sites with census data from 2003-2017, was one of the most complex field studies in Urban Paris, with a dramatic decline of ∼89% of the species over the study period observed [15,16]. In India, a sharp decline in the house sparrow population was observed across Mumbai, Bengaluru, Hyderabad, and other major cities with certain places in India having over 70% decline[17]. Typically the lifespan of the house sparrow is 3 - 5 years in the wild and only about 20 percent of young live past their first year. Cold weather and food availability decide the longevity of Sparrows. *Plasmodium relictum*, a parasite infection, has also affected house sparrow demography across suburban London where sparrows have declined by 71% since 1995[18].

The genome sequencing efforts have been yielding contentious debate heralding short and long read chemistry over the last few years. While the peacock genome has yielded results [19], we have earlier contemplated asking questions on the Passer genome sequencing [20]. The avian genome project[21] [https://b10k.genomics.cn/ last accessed on September 20, 2024] representing several orders is aimed to resolve the phylogeny and collate data for testing hypotheses, understanding extinction and speciation of the birds besides demographic events, their roles of drift and selection in the divergence process [22]. Recently, Magallanes-Alba et al. have done a rapid genome functional annotation pipeline anchored to the house sparrow genome re-annotation and provided transcripts [23]. Our current genome sequencing of house sparrow is based on a muscle of the bird followed by assembly, annotation employing *in silico* approaches using a myriad of tools that could serve as a valuable resource to understand passerines’ evolution, bio/ethno/geography and demography [24].

## Methods

### Sample collection and genome assembly

A bird wing (muscular tissue) was taken from a male house sparrow which was found dead in the lawns of the Zoological Survey of India, Kolkata, and frozen immediately in liquid nitrogen. The sample was handled by Unipath Labs for DNA extraction and sequencing using Illumina HiSeq 2000 platform.The paired end raw reads generated by sequencer were obtained by us for further processing. We performed genome assembly using Shovill (version 1.1.0) /SPAdes (3.15.4) [25]. Initially, we mapped *Gallus gallus* and *Taeniopygia guttata* and the alignment was not successful as upon further alignment using bowtie2, we found 99% mapped reads which led us to conduct downstream scaffolding. The gene completeness for Passer was assessed through Benchmarking Universal Single-Copy Orthologs (Busco version 5.5.0) by using the orthologous genes in the *Gallus gallus* [chicken] genome. We performed a *de novo* assembly rather than a reference-based assembly with a specific set of nucleotide sequences used to represent an organism’s genome [27]. In addition to BUSCO, we employed three tools for assembling the sample, *viz*. Megahit [28], Maryland Super-Read Celera Assembler (MaSuRCA) [29] and Spades. A summary of methods can be found in Figure 1

**Figure 1:**
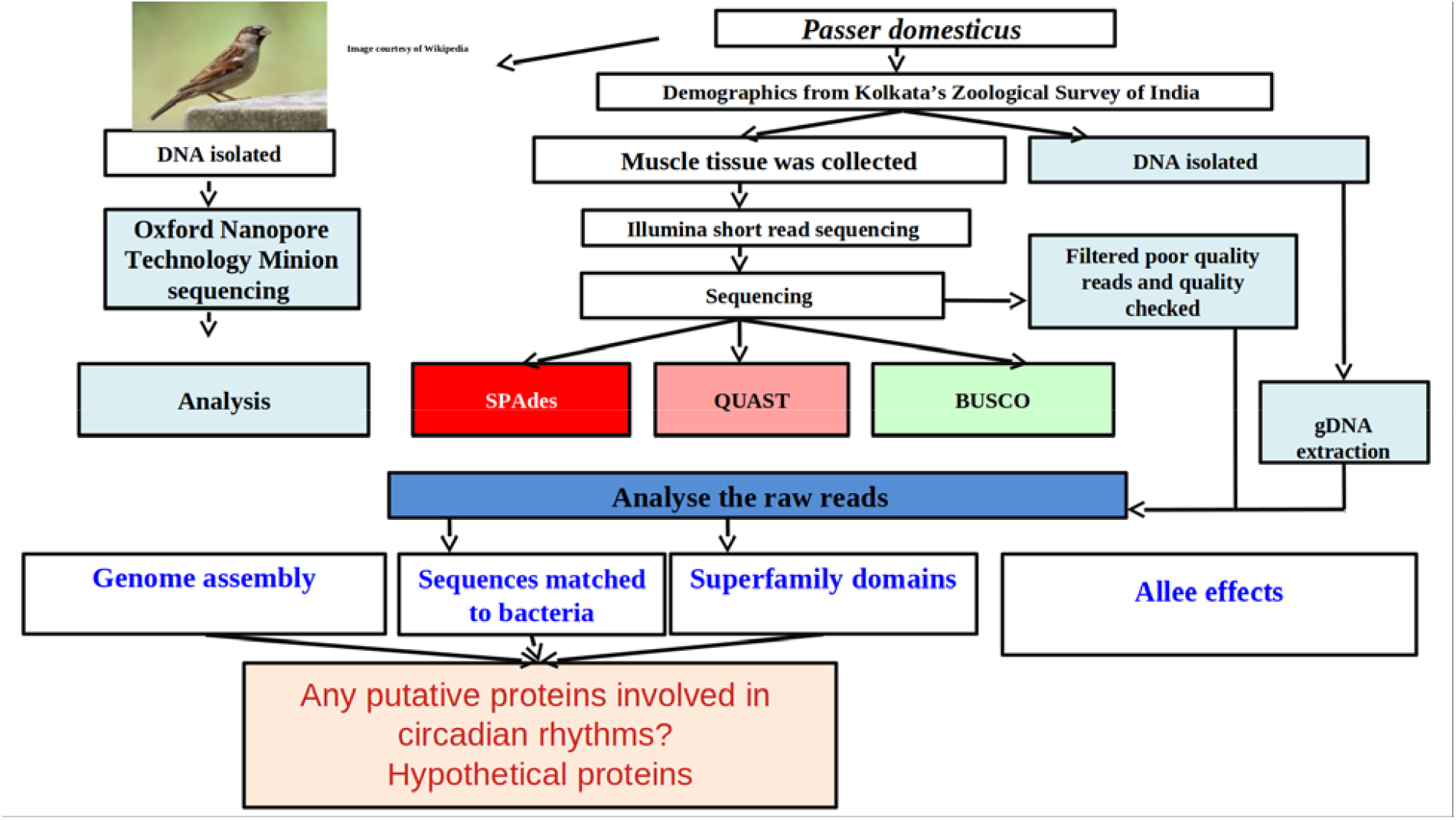
An overview of methodology employed for annotation and assembly of *Passer domesticus*.

### Assembly graph construction

In order to systematically traverse the sequence space and find read overlaps which are crucial for contig assembly, we performed statistical analysis of the assembly to identify the best assembly among the three assemblers as discussed previously, to select the best contigs for further downstream analysis. Statistics of the assembly was done using QUAST to identify the N50 statistics [55]. The ensuing gaps were closed using a command line tool called gap closer, which allowed us to close 3150 gaps constituting the contigs present in our assembly. Prediction of genes in the genome was performed using Augustus [30], an *ab initio* hidden Markov model(HMM)-based gene prediction tool. The scaffolds of the SPAdes were rendered as the input for the Augustus tool against trained species-specific datasets, *viz. Gallus gallus* as a reference for alignment. The obtained GFF file was parsed to get predicted coding sequence and amino acid fasta sequences.

### Mitochondrial genome assembly

We extracted the mitochondrial genome from the reads and the assembly was performed using the GetOrganelle toolkit which includes a number of scripts and libraries[31]. We further employed WGS read data for manipulating and disentangling assembly graphs, and generated reliable organelle genomes, accompanied by labeled assembly graphs and visualized using Bandage[32]. The prediction of the mitochondrial gene from the genome and annotation of the genome was performed using the Mitochondrial genome annotation Server (MITOS2) [33]. It uses BLAST searches with previously annotated protein sequences to predict the protein coding genes and further annotates the tRNAs apart from rRNAs present in the genome. We further performed a BLAST to identify the D-loop, a non-coding region in Mitochondria which acts as the promoter for both light and heavy chains and it is an important feature in mitochondria. We were able to acquire the order of the gene, gene names,start and end of the genes and obtained the values of the intergenic region between genes to construct the mitochondrial DNA. The mitochondrial DNA was constructed and visualized using GenomeVX [34]. For ascertaining codon usage and Ka/Ks, we employed relative synonymous codon usage (RSCU) analysis using Molecular Evolutionary Genetics Analysis Version 11 [35]

### Phylogenetic analysis

The phylogenetic tree construction was performed using MEGA11 followed by reaching consensus with 1Qq tree [36] and MAFFT [37] as we considered species of interest *Passer domesticus: Passer montanus, Passer_domesticus, Passer_ammodendri, Petronia_petronia, Pyrgilauda_blanfordi, Montifringilla_adamsi, Fringilla_polatzeki, Anthus_cervinus, Motacilla_alba, Emberiza_fucata, Spizella_passerina, Agelaius_phoeniceus, Dives_dives, Euphagus_cyanocephalus, Quiscalus_quiscula, Chrysomus_icterocephalus, Pseudoleistes_guirahuro, Molothrus_badius, Gymnomystax_mexicanus, streptopelia_orientalis_voucher_zjbj2* (outgroup). After bootstrapping, we constructed a tree using the unweighted pair group method with arithmetic mean (UPGMA)with a gap penalty of −400 and a gap extension of 0.00. For phylogenetic tree construction, we chose Kimura as the substitution model with bootstrapping set to 100 and the final tree with MAFFT was constructed, While the scoring matrix was given as BLOSUM62 AND 200 PAM, the Jukes-cantor model was set as the substitution model and neighbor joining (NJ) for tree construction wherein bootstrapping was set to 100. For validating the tree inferred to understand the reliability and robustness, we employed a statistical method called the bootstrapping method. From the bootstrapped phylogenetic tree, we concluded that a node is well supported if it remained unchanged after 95 out of 100 iterations of removing one character and resampling our tree; a bootstrap value of 95% indicates this. Using the Mega X tool, we constructed a tree using the UPGMA as an algorithm for the MSA method with a gap penalty of −400 and a gap extension of 0.00. For phylogenetic tree construction, the substitution model chosen was kimura-2 parameter model and method used was maximum likelihood method and bootstrapping values were put to 100 and a tree was obtained. Then finally a tree with MAFFT alignment was constructed, where the MSA and the tree construction was performed with a penalty score rendered as 1.53. The sequences were aligned using progressive method with the tree algorithm as a default parameter.

### Genome annotation, comparison and statistics

Gene annotation was performed with the protein sequence obtained from gene prediction to annotate the genes. Repetitive regions were identified and masked prior to gene prediction using Repeat Modeler [38], a *de novo* transposable element [TE] family identification and modeling package with Repeat Scout tool embedded for identifying the boundaries.[39]. We used the Shovill contigs that were generated as an input with 1391 sequences [1696224 bp] for downstream analyses and the resulting library was later checked with *Gallus gallus* repeat libraries. We compared and checked the annotation using Red [REpeat Detector: (Version 2018.09.10)], a rapid tool for detecting repeats *de novo* on the genomic scale[40].As we searched for prediction of incomplete genes at the sequence boundaries, we also aimed to predict complete genes. The Red GFF file was used as Augustus produced the fasta files containing predicted coding sequences and that were used for Pfamscan searches[Supplementary Table 3a, 3b, 3c) [41]. The resulting Augustus predictions were used as an input with Pfam-A HMM library manually downloaded in stockholm format. We also queried characteristic active site residues if any between the overlaps belonging to the same clans and further checked for functional annotation and domains using InterProScan [42] in inferring the Gene Ontology terms. The *Acanthisitta chloris* genome was obtained from CNGB for comparing the contigs with similarity [43]. Batch Entrez, Smith-Waterman algorithm using UVA FASTA (version 36.3.8i May, 2023) from local searches were made for predicting the proteins [https://fasta.bioch.virginia.edu/fasta/fasta_list.html last accessed on April 1, 2024]. GffCompare was employed to compare predicted sequences at different levels of granularity, thereby annotating the sequences based on their overlaps or proximity to reference annotation transcripts [44]. Sensitivity and precision metrics were computed with GTF file as an input for generating annotated files, yielding the “super-locus” for measuring the accuracy with true positives with other features like true negatives [TN] and false positives [FP] : Sensitivity = TP/[TP+FN] and Precision = TP/[TP+FP]. Finally, RefMap and TMAP files [4] were obtained measuring the reference transcript that either fully or partially matches a transcript from the GTF and those columns in the file describing the most closely matching reference transcript respectively.

### Comparative divergence time estimation

In order to estimate the divergence among selected clade and species, we used BEAST V2.0 [50], where Jmodeltest 2 [51] was used to decide evolutionary model based on BIC value, where best suggested model were HKY I+G. we used mutation rate 0.018 substitution per site per Millions to estimate divergence [52]. The log files were checked using Tracer ensuring that ESS values were more than 200, and final trees were annotated with Treeannotator. Time divergences were estimated using data from all 20 taxa representing seven different families.

## Results and Discussion

### SPAdes achieved better genome assembly statistics when compared to Masurca and Megahit

We obtained the paired end files of 14 GB each which underwent quality check clearance wherein the Phred quality score was improved to 30 after FastP trims which were then assembled. SPAdes achieved better genome assembly statistics when compared to Masurca and Megahit [Table 1]. While the number of contigs were consistent between the Megahit and SPAdes, Masurca yielded a smaller number of contigs as it is not referenced for bird or avian genomes. However, the GC% was found to be consistent with all the three tools attributing to an average of 771 bp. On the other hand, the N50 was comparatively better for SPAdes assembly than the other two assemblers. In summary, *Passer domesticus* genome size was attributed to 922 MB taking the SPAdes assembled genome into consideration.

**Table 1:** Assembly features extracted from three different tools, *viz*. Megahit, SPAdes and Masurca. From all the results the one with better N50 and avg value was the SPAdes assembly.

| Tools>>> | Megahit | SPAdes | MaSuRCA |
| --- | --- | --- | --- |
| contigs (N) | 1018705 | 1000366 | 608253 |
| Length | 866 MB | 922 MB | 830 MB |
| Largest contig (BP) | 83946 | 87277 | 78946 |
| N50 (BP) | 1027 | 1343 | 740 |
| N90 (BP) | 434 | 399 | 400 |
| **N's (BP)** | 0.00 | 0.00 | 0.00 |
| **GC %** | 41.74 | 41.93 | 41.49% |

### Gene prediction yielded 24152 genes across as many as 45634 transcripts

When genes were predicted using Augustus, and mapped from the BWA-mem of *Gallus gallus*, a considerable similarity of ca. 80.3 % was achieved further attributing to 24152 genes, 38972 introns and 45634 transcripts. The GFF file produced was then parsed to check downstream functional annotation tools using the KOG, non-redundant (NR) and Uniprot databases. The consensus hits were searched with “uniq sort” to find the most occurring species and considered the 10 best hits. We observed that NR yielded better results with *Passer montanus* occurring the most with a count of 7308, 1365 *Stutzerimonas stutzeri*, 939 *Melospiza melodia maxima*, 788 *Pyrgilauda ruficollis*, 780 *Hirundo rustica rustica*, 715 *Chloebia gouldiae*, 679 *Limosa lapponica baueri*, 652 *Onychostruthus taczazanowskii*, 457 *Lonchura striata domestica* and 319 *Motacilla alba*. What remains intriguing was that we detected the presence of bacterial sequences among the hits which was quite surprising. We argue that this could be due to the contamination caused during sequencing or any horizontal gene transfer events that might have happened over the years or birds could include a lot of symbiotic or pathogenic bacteria which can be part of their microbiota. We also observed the presence of many hypothetical proteins indicating that they are known unknown regions implying the need to annotate the genome significantly. There were the presence of many genes which are important for adaptations such as vocal learning, circadian rhythms, allee effects etc. BMAL1, clock genes were found to be associated in our list which are known to maintain the circadian rhythms of the species. We also found TLR4 which helps in identifying pathogens and initiation of immune responses besides HBAA, HBB, HBA genes that are known for oxygen transport which helps in adaptation.

### Mitochondrial genome yielded a genome of size 16,804 bp

Gene prediction and annotation predicted a total of 37 genes from which 13 genes were protein coding genes and 22 were tRNAs, 2 rRNAs and the non-coding region which is the D-loop was also identified.A labeled mitogenome was produced using GenomeVX online platform (Figure 2). The relative synonymous codon usage (RSCU) analysis of the codons was calculated in which the lowest preferences are AGA (R), AGG(R), ACG (T) [Table 2]. While they are less frequently used due to the lower level of those codon specific tRNAS, they might slow down the translation of proteins efficiently thus avoiding the use of these codons in efficient protein translation. We observed different patterns of selective pressure by analyzing the Ka/Ks ratios for the specified genes with strong purifying selection seen in the majority of the genes, which have Ka/Ks ratios much less than 1, including cox1 (0.003), Cox3 (0.012), Nad2 (0.017), Nad3 (0.02), cob (0.02), NAD1 (0.034), Nad4l (0.035), Nad5 (0.03), and Nad6 (0.048). Given that the proteins these genes encode are essential to the organism’s survival and functionality and that harmful mutations are progressively eliminated, it is likely that these genes are highly conserved. The Atp6 gene, on the other hand, appears to be under positive selection, indicating that natural selection may be favoring modifications in the protein product of this gene that may be advantageous with a Ka/Ks ratio of 1.03 [Table 3; Figure 2]. While the scoring matrix was given as BLOSUM62 AND 200 PAM, the Jukes-cantor model was set as the substitution model and NJ for tree construction wherein bootstrapping was set to 100 (Figure 3).

**Table 2:** The RSCU values provided for the mitochondrial genome with the codons CUA (L), GCC(A), CGA(R) having highest values indicate more frequent occurrence. They might be preferred due to the abundance of tRNA contributing to efficient protein translation and protein stability reflecting evolutionary adaptation for protein synthesis. The codons that are translated efficiently are preferred in highly expressed genes

| Codon | RSCU | Codons | RSCU | Codons | RSCU | Codons | RSCU |
| --- | --- | --- | --- | --- | --- | --- | --- |
| UUU(F) | 0.27 | UCU(S) | 0.53 | UAU(Y) | 0.44 | UGU(C) | 0.26 |
| UUC(F) | 1.73 | UCC(S) | 2.08 | UAC(Y) | 1.56 | UGC(C) | 1.74 |
| UUA(L) | 0.47 | UCA(S) | 1.83 | UAA(*) | 0.28 | UGA(*) | 2.69 |
| UUG(L) | 0.18 | UCG(S) | 0.35 | UAG(*) | 0.03 | UGG(W) | 1 |
| CUU(L) | 0.34 | CCU(P) | 0.45 | CAU(H) | 0.32 | CGU(R) | 0.49 |
| CUC(L) | 1.23 | CCC(P) | 1.42 | CAC(H) | 1.68 | CGC(R) | 1.15 |
| CUA(L) | 3.08 | CCA(P) | 1.84 | CAA(Q) | 1.64 | CGA(R) | 3.7 |
| CUG(L) | 0.71 | CCG(P) | 0.29 | CAG(Q) | 0.36 | CGG(R) | 0.49 |
| AUU(I) | 0.52 | ACU(T) | 0.58 | AAU(N) | 0.31 | AGU(S) | 0.14 |
| AUC(I) | 1.61 | ACC(T) | 1.71 | AAC(N) | 1.69 | AGC(S) | 1.07 |
| AUA(I) | 0.86 | ACA(T) | 1.67 | AAA(K) | 1.7 | AGA(R) | 0.08 |
| AUG(M) | 1 | ACG(T) | 0.04 | AAG(K) | 0.3 | AGG(R) | 0.08 |
| GUU(V) | 0.83 | GCU(A) | 0.48 | GAU(D) | 0.29 | GGU(G) | 0.38 |
| GUC(V) | 0.93 | GCC(A) | 2.22 | GAC(D) | 1.71 | GGC(G) | 0.95 |
| GUA(V) | 1.64 | GCA(A) | 1.19 | GAA(E) | 1.47 | GGA(G) | 1.9 |
| GUG(V) | 0.61 | GCG(A) | 0.11 | GAG(E) | 0.53 | GGG(G) | 0.77 |

**Table 3:**
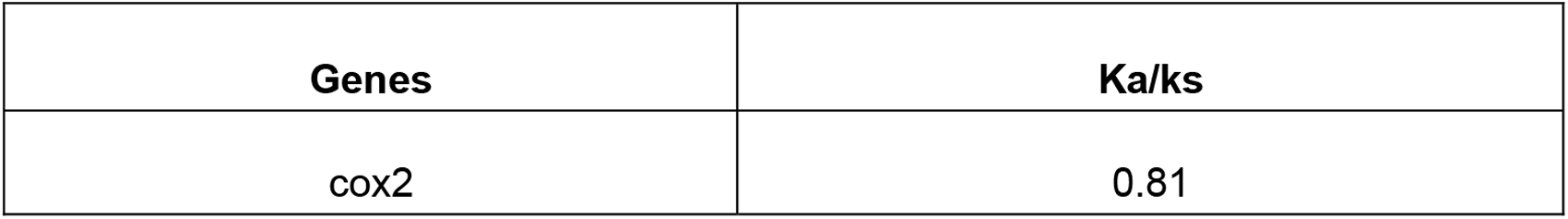
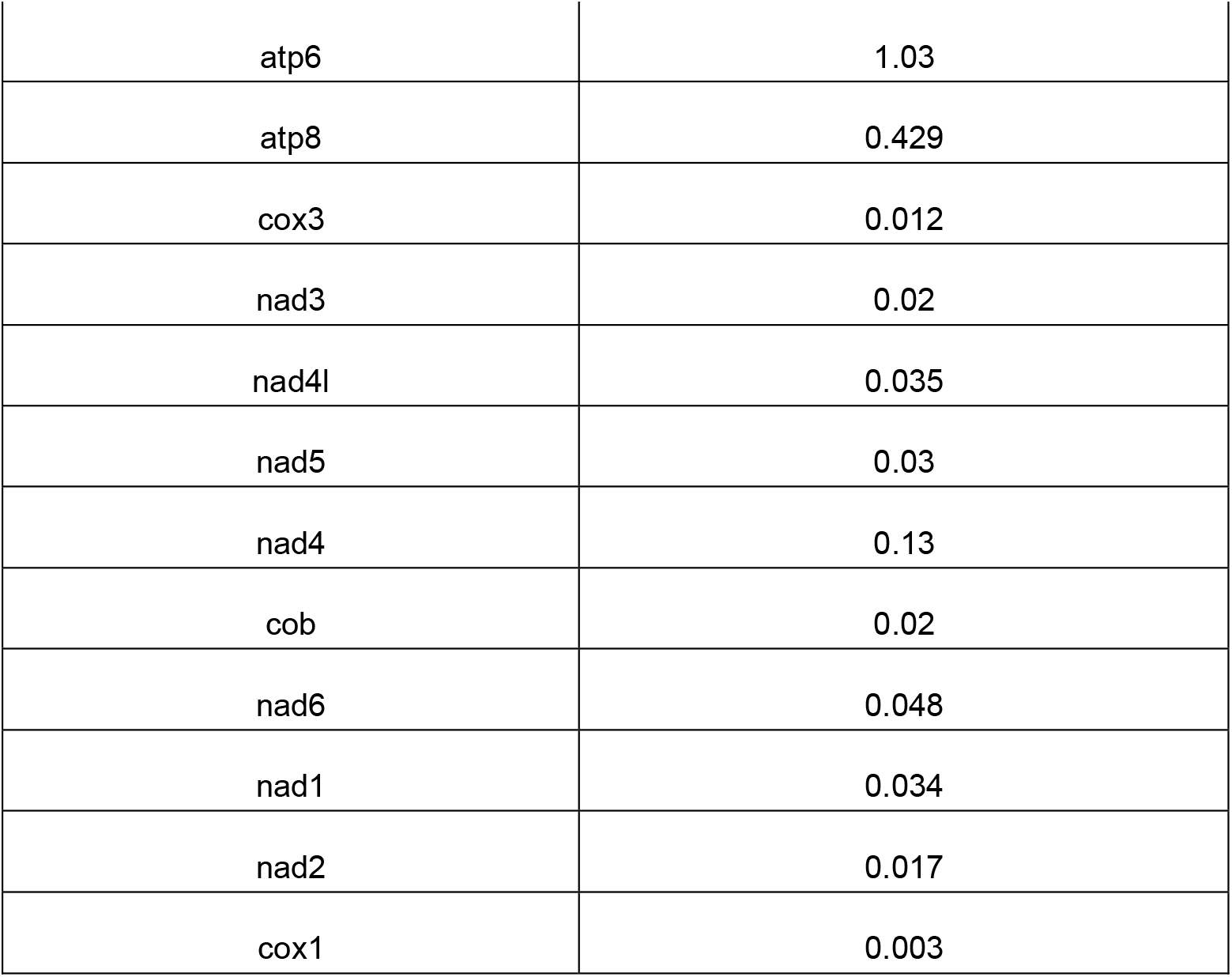
ka/ks values indicating all the genes except atp6 have values less than one indicating they are undergoing strong purifying selection whereas the atp6 undergoes a positive selection with a value greater than one.

**Figure 2:**
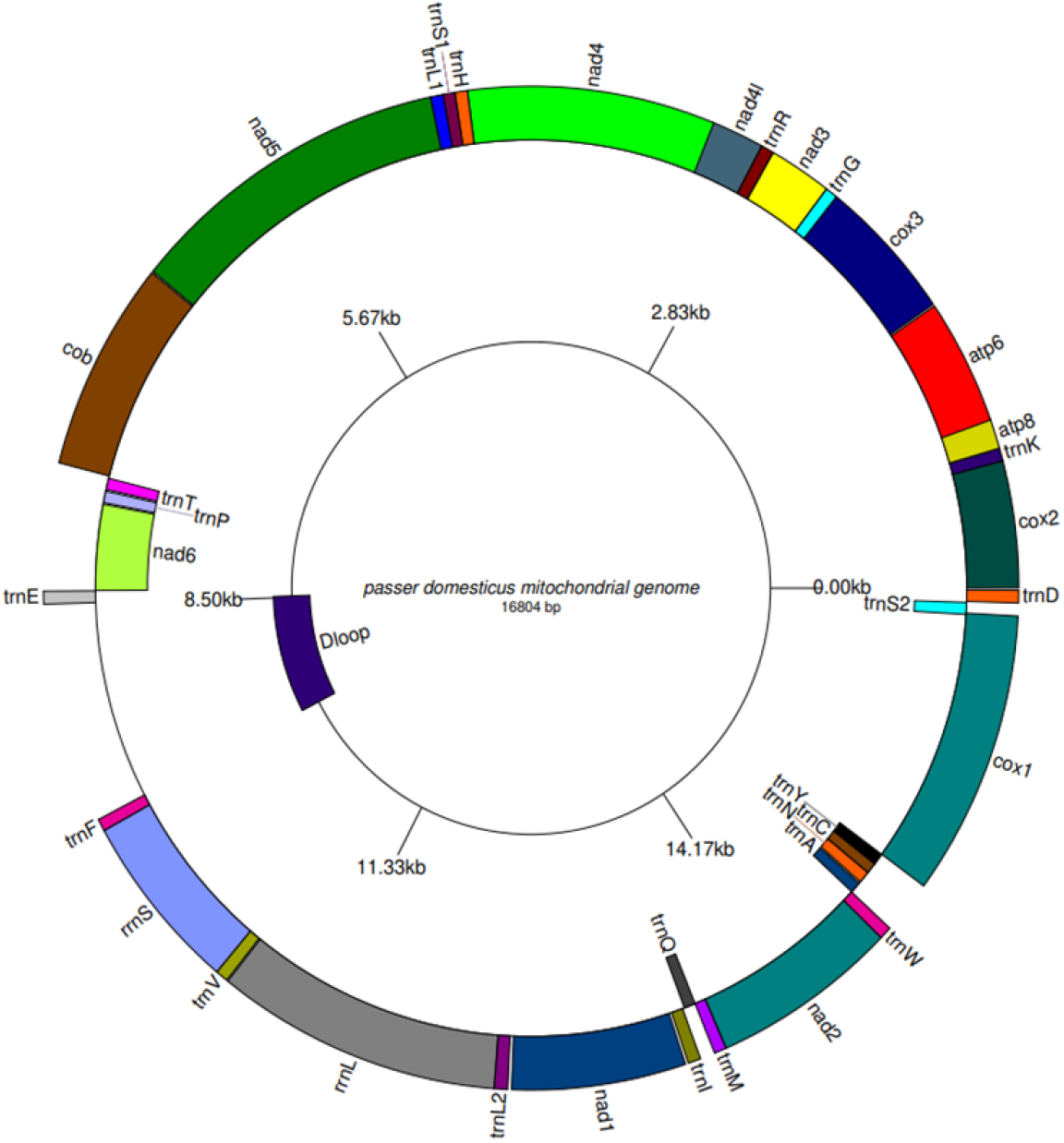
Labeled mitogenome representing the 37 genes including the protein coding gene tRNAs,rRNAs and the non-coding region, which is the control region or the d-loop

**Figure 3.**
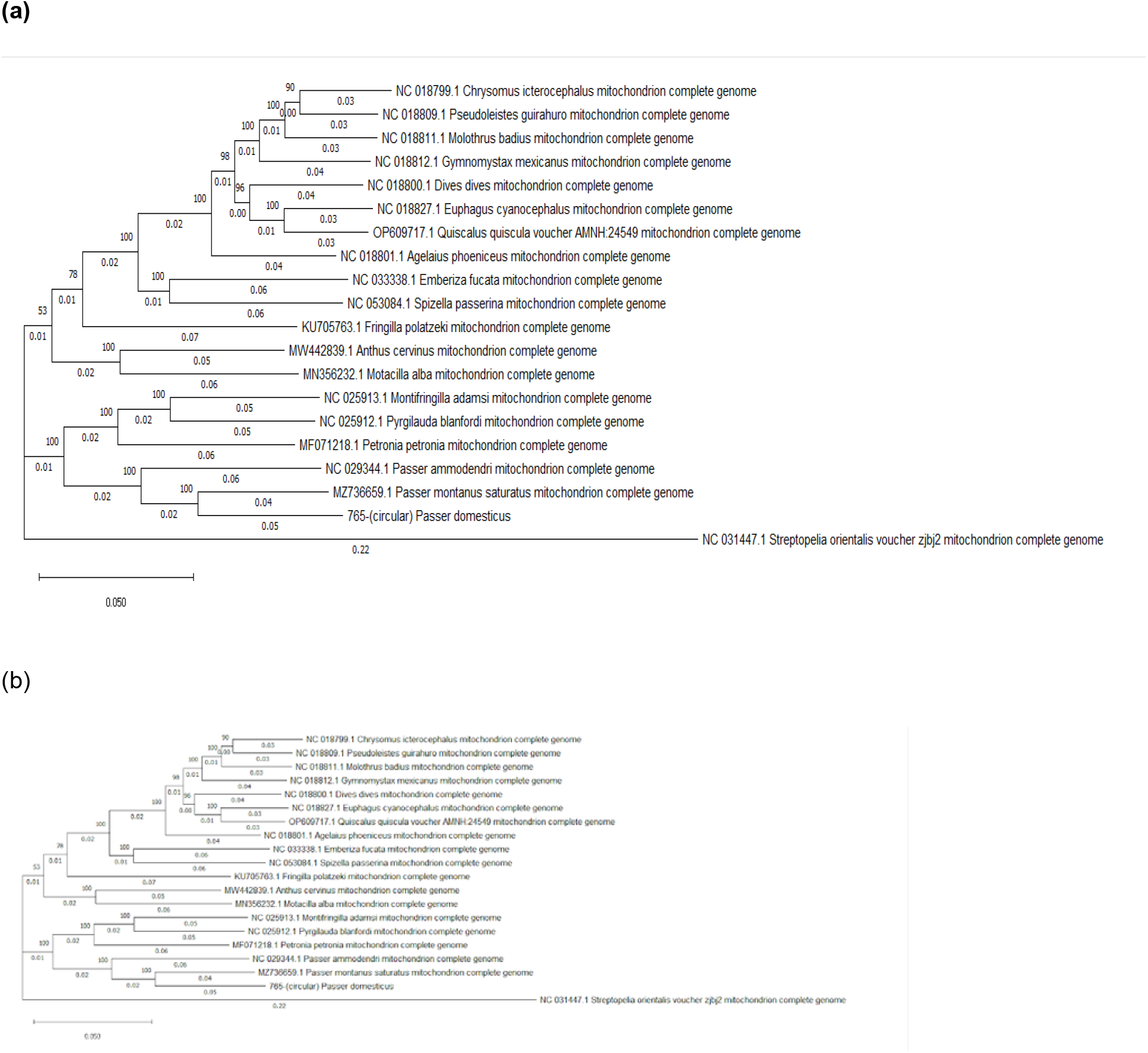
(a): Phylogenetic tree representing all the 20 species, in which the *Passer domesticus* is closely related to including *Passer montanus, Passer ammodendri, Petronia petronia, Pyrigilauda blandford*i and *Montifringilla adamsi* having highly supported bootstrap values ie,100. Whereas less closely related the other species such as *Motacilla alba, Anthus cervinus* all belong to the branches with bootstrapping value of 78, which is poor. The *streptopelia_orientalis_voucher_zjbj2* shows the most distant species in the tree which is the outgroup species with a low supported bootstrap value 53. (b)

Avian genomes are ca. 70% smaller than mammalian genomes and Passerines being small are thought to maintain the contiguity that we have shown as compared to other vertebrates. We are limited by the amount of genomic attributes coming from non-coding regions, nevertheless, we argue that this could be because of the large number of hypothetical proteins which demand accurate annotation. While many genomes show synteny, we are planning to sequence more genomes that could provide us insights on adaptation, evolutionary mechanisms, substitutions, and importantly the circadian rhythms or electromagnetic effects that these birds might be associated with, thus drifting away from the urban life.

The Passer genome showed considerable similarity with bacterial sequences indicating that a large number of plausibly gut microbiota may have evented horizontal gene transfer [HGT] over the years [45]. This could be attributed to the bird’s foraging on insects and small grains with bacteria inhabited in them could exist with endosymbionts and their invertebrate hosts, particularly insects. We argue that these could in principle be largely associated with modern evolutionary theory promoting such gene transfer. In addition, the alignment between these genomes is indicative of inversions/transversions which is in agreement with bird genomes [46]. The horizontal gene transfer has been described between bacteria and animals, and such transfers have potential to influence evolution of animals as well. Although HGT is less common among Eukaryotes, however has been discovered in different taxa of Eukaryotes, i,e., from Drosophila melanogaster to Drosophila willistoni (Daniels et al., 1990), among fishes (Graham et al., 2008), from Fungi to pea aphids (Moran and Jarvik, 2010), from Trypanosoma cruzi to Human (Hecht et al., 2010) The 710 contigs were mapped to the GenBank yielding 152 hypothetical proteins indicating that a large amount of proteins emerged from the known-unknown regions, implying the need to annotate the genome significantly {Supplementary Table 1). Among these, 252 sequences from protein domains match to animals while the rest match to bacteria. We observed a large number of Kelch domains in addition to a PHD finger 1 found in Histone-lysine N-methyltransferase 2C (KMT2C) and 2D (KMT2D) protein that is largely conserved. Also termed myeloid/lymphoid or mixed-lineage leukemia protein 3 (MLL3), it is associated with circadian factors contributing to genome-scale circadian transcription. These observations could indicate that a large number of genes possibly have undergone divergent evolution or the genes might have been lost and found through speciation. This fragmented genome is also indicative of BUSCO statistics from avian genomes that were reported earlier in Chaffinch genomes [47].

Our annotation level of sequence representation is scanty but diverse as we found increased gene counts in comparison to *Gallus gallus* [53]. However, a large fraction of our genes remain unmapped and undiscovered in the Chicken genome. We deem that ours is a draft genome, nonetheless, small species such as Passerines still have no distinct function diversity and their demand for evidence can only be classified through experimentation. While we attempted to create a good quality *de novo* based reference genome, the final assembly encompassing 922 MB is in agreement with the Elgvin et al. [54] works of a similar sized genome albeit the medium-density linkage map and order, the assembly scaffolds into chromosomes was meticulously done across several samples which in our case we couldn’t as the sample acquisition has been a major limitation. Nevertheless, the genes were mapped to their reference genome for downstream analysis [Table 4].

**Table 4:** comparison of genes that were mapped to the reference genome.

| S.No | Gene Name | NCBI accession | Contigs mapped |
| --- | --- | --- | --- |
| 1 | APC | NC_087512.1:52445657-52540315 | NODE_14189_length_3730_cov_2.598686 |
| 2 | HSDL2 | NC_087512.1:69765571-69788616 | NODE_20005_length_3394_cov_4.411215 |
| 3 | ETFA | NC_087487.1:960048-989096 | NODE_45445_length_2642_cov_2.323977 |
| 4 | NOTCH2 | NC_087480.1:32766603-32849060 | NODE_2951_length_5290_cov_3.101669 |
| 5 | NOTCH1 | NC_087491.1:10550409-10592339 | NODE_82034_length_2127_cov_3.214634 |
| 6 | GLI3 | NC_087474.1:51942133-52148229 | NODE_5580_length_4619_cov_3.460810 |
| 7 | WNT4 | NC_087495.1:277607-294952 | NODE_126693_length_1756_cov_2.892198 |
| 8 | TIMP2 | NC_087493.1:1552785-1561149 | NODE_92905_length_2020_cov_2.762738 |
| 9 | RBP1 | NC_087484.1:25002122-25019513 | NODE_19406_length_3423_cov_2.725344 |
| 10 | PRK11A | NC_087512.1:13609738-13635725 | NODE_48977_length_2574_cov_3.450941 |
| 11 | GNAI1 | NC_087478.1:9772700-9805360 | NODE_5061_length_4724_cov_3.114698 |
| 12 | CRHR1 | NC_087500.1:437197-467780 | NODE_32621_length_2944_cov_3.164632 |
| 13 | ZFAND5 | NC_087512.1:62564205-62578406 | NODE_86240_length_2083_cov_4.130608 |
| 14 | SECISBP2 | NC_087512.1:42294093-42319272 | NODE_2582_length_5428_cov_3.587927 |
| 15 | REEP5 | NC_087512.1:52407038-52426127 | NODE_48707_length_2579_cov_3.693046 |
| 16 | MIA3 | NC_087476.1:85960932-85989020 | NODE_12474_length_3848_cov_3.843543 |
| 17 | A2M | NC_087475.1:87627899-87657608 | NODE_583_length_7451_cov_3.871576 |
| 18 | KCNN2 | NC_087512.1:68344752-68419555 | NODE_16813_length_3560_cov_3.043928 |
| 19 | BNC2 | NC_087512.1:33461143-33796687 | NODE_5333_length_4668_cov_3.558048 |

### Seven superfamily domains are conserved between *Passer domesticus and Acanthisitta chloris*

The final set of contigs obtained for assembly were mapped against NCBI taxa [40] even as it matches to 14947 contigs with 769 of them making 100% query coverage with various hits from taxa. Amongst the superfamily members, intermediate filament protein, BASP1: Brain acid soluble protein 1, glycine rich LPXTG-anchored collagen-like adhesin and collagen with keratin and some hypothetical protein-FTsK translocase domains. What remains intriguing is the presence of probable chromatin-remodeling complex ATPase chain which is possibly associated with Allee effects that occur when individual fitness suffers at low population size or density [48]. We also argue that the genetic architecture and community-wide admixture could provide insights into their evolution [49]. On the other hand, keeping in view of the sample characteristics as a measure of genome completeness, we also mined the long read data from Oxford nanopore technologies [data not shown]. While the aforementioned statistics in the form of N50, N90, BUSCO are optimized for short reads, we were short of full potential in analyzing the genomic reads generated from long reads. Furthermore, the sequences generated need to be polished to overcome the mis-assemblies and have accurate scaffolding for generating more suitable short reads. On divergence, Columbidae appeared to have diverged early compared to others and was considered an outgroup in this analysis. In our sequenced sample, the house sparrow belonged to the family Passeridae. The divergence and expansion of major families began around 7-9 million years ago. Our studied sample formed a subclade with the Eurasian tree sparrow (*P. m. saturatus*) and Saxual sparrow (*P. ammodendri*), having a Time to Most Recent Common Ancestor (TMRCA) of 4.4 million years ago. Furthermore, our analysis yielded a TMRCA of 2.9 million years ago for *Passer domesticus* and *P. m. saturates* (Figure 4).

**Figure 4:**
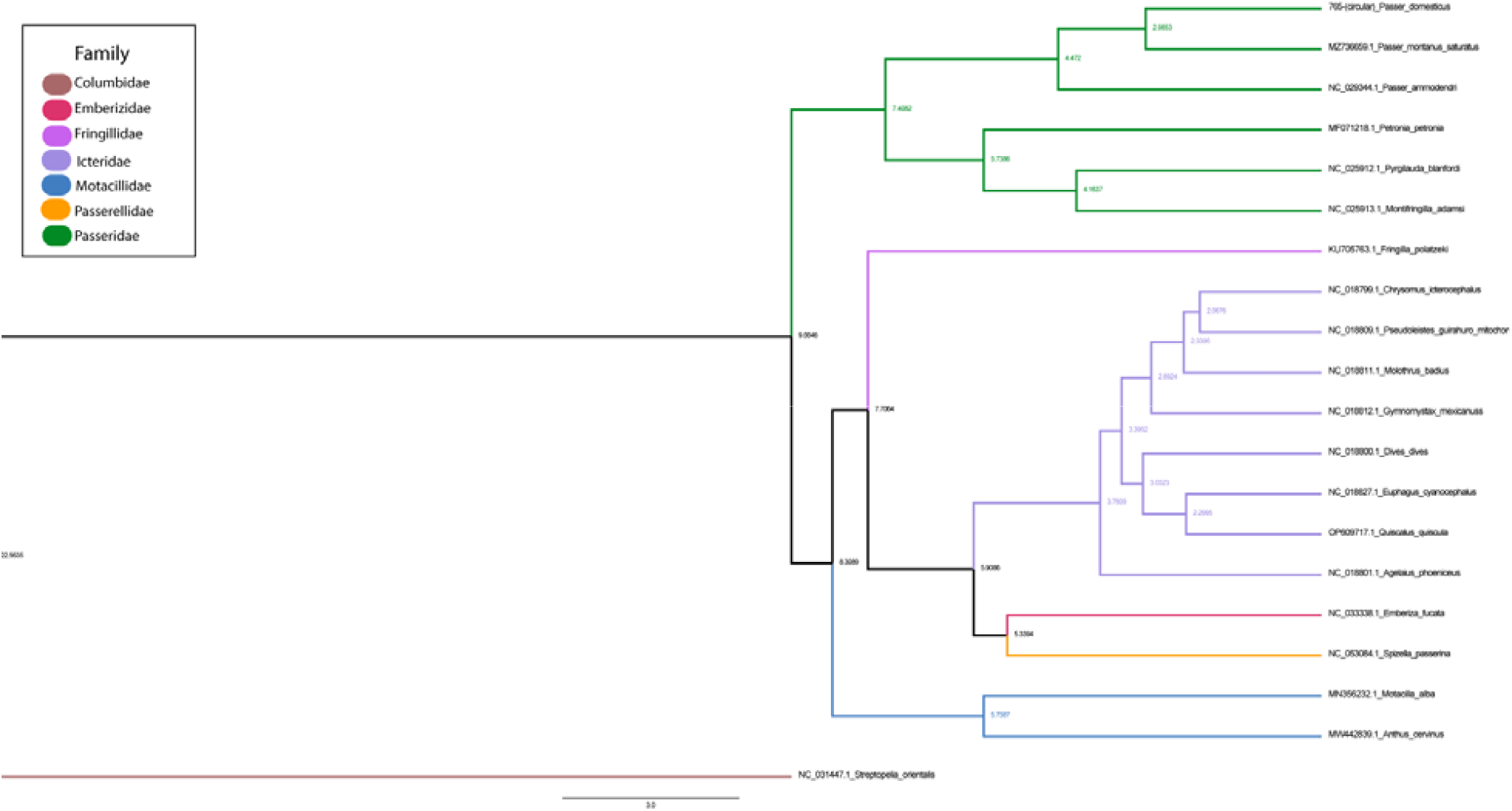
A time-calibrated tree with 20 selected taxa using complete mitochondrial DNA sequences The tree was generated using Beast v2.5 with 20 million MCMC runs. Each respective family is represented by a different color, and the node labels indicate the age in million years.

## Conclusions

We provide here a high-quality whole genome sequencing and assembly for the house sparrow, *Passer domesticus* which could serve us to ask a large number of questions from genomics. Our annotation serves as a useful resource to check adaptation, divergence and speciation. The orthologous annotation and protein mapping, bacterial correlates show that the HGT events may not be ruled out.

## Acknowledgments

PS gratefully acknowledges Department of Forestry, Governments of Rajasthan and India, Zoological Survey of India, and Birla Institute of Scientific Research for providing the apriori permission for collection of bird’s sample.

## Authors’ contributions

PS ideated the project with GC and PBK. VK,SS, SP, AT, GS, DV and PS contributed equally to the work. All the other authors worked towards improvement of tables and figures. PS, GC and PBK proofread the manuscript.

## Data Availability

The genome assembly has been deposited at NCBI under BioProject ID: PRJNA1027087.

